# Genome-wide association reveals host-specific genomic traits in *Escherichia coli*

**DOI:** 10.1101/2022.02.08.479532

**Authors:** Sumeet K. Tiwari, Boas C.L. van der Putten, Thilo M. Fuchs, Trung N. Vinh, Martin Bootsma, Rik Oldenkamp, Roberto La Ragione, Sebastien Matamoros, Ngo T. Hoa, Christian Berens, Joy Leng, Julio Álvarez, Marta Ferrandis-Vila, Jenny M. Ritchie, Angelika Fruth, Stefan Schwarz, Lucas Domínguez, María Ugarte-Ruiz, Astrid Bethe, Charlotte Huber, Vanessa Johanns, Ivonne Stamm, Lothar H. Wieler, Christa Ewers, Amanda Fivian-Hughes, Herbert Schmidt, Christian Menge, Torsten Semmler, Constance Schultsz

## Abstract

*Escherichia coli* is an opportunistic pathogen that can colonize or infect various host species. There is a significant gap in our understanding to what extent genetic lineages of *E. coli* are adapted or restricted to specific hosts. In addition, genomic determinants underlying such host specificity are unknown.By analyzing a randomly sampled collection of 1198 whole-genome sequenced *E. coli* isolates from four countries (Germany, UK, Spain, and Vietnam), obtained from five host species (human, pig, cattle, chicken, and wild boar) over 16 years, from both healthy and diseased hosts, we demonstrate that certain lineages of *E. coli* are frequently detected in specific hosts. We report a novel *nan* gene cluster, designated *nan-9,* putatively encoding acetylesterases and determinants of uptake and metabolism of sialic acid, to be associated with the human host as identified through genome wide association studies. *In silico* characterization predicts *nan-9* to be involved in sialic acid (Sia) metabolism. *In vitro* growth experiments with a representative Δ*nan E. coli* mutant strain, using sialic acids 5-*N*-acetyl neuraminic acid (Neu5Ac) and *N*-glycolyl neuraminic acid (Neu5Gc) as the sole carbon source, indicate an impaired growth behaviour compared to the wild-type. In addition, we identified several additional *E. coli* genes that are potentially associated with adaptation to human, cattle and chicken hosts, but not for the pig host. Collectively, this study provides an extensive overview of genetic determinants which may mediate host specificity in *E. coli.* Our findings should inform risk analysis and epidemiological monitoring of (antimicrobial resistant) *E. coli.*

## Introduction

*Escherichia coli* is a Gram-negative bacterium which has been isolated from various host species, including humans, cattle, chickens and pigs(1). Because *E. coli* can colonize or infect multiple host species, this bacterium can act as a reservoir for genes encoding antimicrobial resistance (AMR)(2) that can be transmitted between different host species. The likelihood that *E. coli* and its AMR encoding genes persist in a new host after transmission depends on multiple factors(3,4). For example, small changes in metabolic pathways may enable *E. coli* to colonize or infect a host more efficiently(1). Several studies have suggested that highly successful *E. coli* clones, such as the sequence type 131 (ST131) clone(5,6) or clonal complex 87 (ST58 and ST155) *E. coli* facilitate the spread of AMR *E. coli* in the human population(7) whilst other studies have shown that different lineages of AMR *E. coli* vary in their ability to spread(8). These findings both indicate that AMR genes, at least to some extent, hitchhike on bacterial strains that are specifically equipped to colonize a given host. Beyond classical virulence or adhesion factors, genetic and functional traits defining different degrees of host adaptation(3,9) and thereby indirectly impacting on the spread of AMR between host species, have not been identified thus far.

Comparative genomic analysis of bacterial populations from multiple hosts has revealed signatures of host-adaptation in bacterial genomes(10). The emergence of large-scale bacterial genome-wide association studies (GWAS) allowed for the detection of genes or genomic variants that are associated with resistance, pathogenicity, and host adaptive traits(11–13). Here, we have applied population-based bacterial GWAS to identify host-associated genomic determinants in a diverse panel of 1,198 *E. coli* isolates, irrespective of their AMR pattern. Isolates were recovered from five different host species, including healthy and diseased individuals from four different countries in two continents over 16 years. The *pan-genome* was analyzed for specific host association followed by a *k-mer* based bacterial GWAS approach to identify host-specific genomic determinants and their potential role in host-adaptation.

## Material and Methods

### a) Sampling strategy

A panel of 1213 *E. coli* isolates from four countries (Germany, UK, Spain, and Vietnam), obtained from five host species (human, pig, cattle, chicken, and wild boar) during three time periods (2003-2007, 2008-2012 and 2013-2018) from both healthy and diseased hosts were selected randomly from existing strain collections and newly collected isolates. Out of 120 possible strata (defined as a unique combination of country, host, time-period, and host health status), 42 strata contained isolates. We included all isolates available per stratum if there were less than 30 isolates and performed a random selection of up to a maximum of 30 isolates if more were available. Potentially duplicate isolates that were part of an outbreak, isolated at a single location within a short timeframe, or from a single farm or a single individual were excluded. Only one isolate per individual was included in the analyses. Isolates included per stratum are shown in Table S1.

### b) DNA extraction and sequencing

The DNA of the *E. coli* isolates from Germany was extracted using the QIAamp DNA Mini Kit (Qiagen) following the manufacturer’s instructions. The DNA concentration was evaluated fluorometrically by using Qubit™ 2.0 fluorometer (Invitrogen, USA) and the associated Qubit™ dsDNA HS Assay Kit (0.2-100ng) and Qubit™ BR Assay Kit (2-1000ng), respectively. The libraries were generated using Nextera DNA library preparation (Illumina, https://www.illumina.com). The sequencing was performed using the Illumina MiSeq and HiSeq systems, generating 2 × 250 bp and 2 × 150 bp reads, respectively.

The DNA of the *E. coli* isolates from the UK was purified using a Promega DNA Wizard^®^ genomic purification kit and quantified using Nanodrop. Libraries were generated using Nextera XT technology (Illumina), and DNA sequencing of isolates was performed at the Animal and Plant Health Agency (APHA, Surrey, UK, https://www.gov.uk/government/-organisations/animal-and-plant-healthagency) using an Illumina MiSeq system generating 2 × 150 bp reads.

For *E. coli* isolates from Spain, DNA was extracted using the DNA blood and tissue Qiagen kit according to the manufacturer’s instruction. The total amount of DNA was quantified using a Qubit fluorometer and frozen at −20°C until further analysis. Libraries were prepared using Nextera XT DNA Library preparation (Illumina), and DNA samples were sequenced using a MiSeq platform (2 × 300 cycle V3 Kit).

The DNA of the *E. coli* isolates from Vietnam was extracted using the Wizard Genomic DNA purification kit (Promega, Madison, WI, USA) following the manufacturer’s instructions. The concentration of the DNA was measured fluorometrically by using picogreen (Invitrogen). The sequencing was performed using an Illumina HiSeq 4000 system, which generates 2 × 150 bp reads.

### c) Quality control

Adapter sequences were removed from raw reads using flexbar v3.0.3(14,15) with trimming mode (-ae) ANY. Low-quality bases within raw reads (Phred score value <20) were trimmed using a sliding window approach (-q WIN). FastQC v0.11.7(16) and MultiQC v1.6(17) were used for quality control before and after processing steps.

### d) Genome assembly and annotation

Adapter-trimmed reads were assembled using SPAdes v3.13.1(18) using read correction. Scaffolds smaller than 500bp were discarded. QUAST v5.0.0(19) was used to assess assembly quality using default parameters. Draft assemblies were excluded if the N50 was below an aribrary value of 30 kbp or consisted of more than 900 contigs. Draft genomes were annotated using prokka v1.13(20) with a genus-specific blast for *Escherichia.* Phylogroups were predicted using ClermonTyper v1.4.1(21), and sequence types (STs) of the isolates were identified *in silico* using the Achtman seven gene MLST scheme using mlst (https://github.com/tseemann/mlst).

### e) Pan-genome and phylogenetic analysis

Roary v3.12.0(22) was used to define the *pan-genome* of the population, using paralog splitting. The core genes were aligned using prank(23) on default parameters. The core gene alignment was used to construct the phylogenetic tree using RaxML 8.2.4(24) with 100 bootstraps under a General Time Reversible (GTR) substitution model with the Gamma model of rate heterogeneity and Lewis ascertainment bias correction(25). The core gene phylogeny was corrected for recombination using ClonalFrameML(26) using default parameters. Phylogenetic Clusters (or BAPS clusters) within the dataset were defined using hierBAPS(27,28) based on the core gene alignment. The accessory gene clustering was performed using package Rtsne v0.15(29,30) with 5000 iterations and perplexity 15 in R v3.6.1. iTOL(31) and Microreact(32) were used to visualize the population structure in the context of available metadata. The function chisq.test from the MASS library(33) (v7.3-51.1) was used in R(34) (v3.5.2) to perform X^2^-tests of independence between phylogenetic clusters and host species. Tests were carried out on the full dataset (14 phylogenetic clusters vs. five hosts and nine phylogroups vs. five host species).

### f) Genome-wide association study (GWAS)

We excluded the wild boar *E. coli* isolates from the GWAS analysis, because of their low number (n=29). GWAS was performed to screen *k-mers* for associations with their host (pig, human, chicken, and cattle). Assemblies were shredded into *k-mers* of 9-100 bases using FSM-lite (https://github.com/nvalimak/fsm-lite). The association between *k-mers* and host phenotype was carried out using Fast-LMM linear mixed model implemented in pyseer(35) using a pairwise similarity matrix derived from the phylogenetic tree as population correction. A GWAS analysis was carried out for each host (pig, human, chicken, and cattle). To reduce false-positive associations, isolates from the host of interest were compared with an equal number of isolates from each of the other hosts, designated control isolates. This analysis was repeated 100 times per host of interest by selecting the control strains from other hosts per iteration(36). The selection of control isolates was random and with replacement except for stratification by phylogenetic clusters to minimize phylogenetic bias. The statistical significance threshold was estimated based on the number of unique *k-mers* patterns for each run(35). *K-mers,* which were significantly associated with 90% of the runs per host, were retained and mapped to reference genomes (Table S2) using a fastmap algorithm in bwa(35,37). An arbitrary cut-off of a minimum of 10 *k-mers* mapped per gene was chosen for further analysis to reduce false-positives. *In silico* characterization and gene ontology (GO) assignment was performed using Blast2GO(38), and Clusters of Orthologous Groups (COGs) were assigned using CD-search(39,40).

### g) Prevalence of a human-associated *nan* gene cluster

All available *E. coli* genome assemblies in NCBI RefSeq were downloaded on Nov 29^th^, 2019, using NCBI-genome-download (https://github.com/kblin/ncbi-genome-download). Using a custom ABRicate (https://github.com/tseemann/abricate) database, consisting of the nine genes of the novel human-associated *nan* gene cluster, all downloaded genomes (n=17994) were scanned. STs for all the genomes were assigned as described above.

### h) Construction of mutants and phenotypic experiments

Mutants *Δnan-9* (Amp^R^) and Δ*nanRATEK* of extra-intestinal pathogenic *E. coli* (ExPEC) strain IMT12185 (ST131; RKI 20-00501; Amp^R^) were constructed using the Datsenko-Wanner method(41). The genomic DNA of the wild-type and the mutant strains was isolated using a QIAamp DNA Mini Kit (QIAGEN). Libraries were prepared using the Nextera XT DNA Library preparation kit (Illumina), and MinION one-dimensional (1D) libraries were constructed using the SQK-RBK004 kit (Nanopore technologies, Oxford, UK) and loaded according to the manufacturer’s instructions onto an R9.4 flow cell. MinIon sequencing data were collected for 48 h and the paired-end Illumina sequencing was performed using MiSeq. Hybrid assembly using Illumina and MinION reads was performed using unicycler v0.4.8(42) with default parameters to complete both strains’ genomes. The absence of the desired genes was confirmed based on the assembly followed by annotation using prokka v1.13(20).

Carbon utilization and chemical sensitivity of the deletion mutants and their parental strain were tested using a Biolog Phenotypic Array system, using the PM1 MicroPlate and the Gen III MicroPlate according to the manufacturer’s instructions.

### i) Growth curve analysis

*E. coli* strains were grown at 37°C aerobically in lysogeny broth (LB) (10 g/l tryptone, 5 g/l yeast extract, 5 g/l NaCl, pH 7.5) or in minimal medium (MM). MM is M9 mineral medium (33.7 mM Na_2_HPO_4_, 22.0 mM KH_2_PO_4_, 8.55 mM NaCl, 9.35 mM NH_4_Cl) supplemented with 2 mM MgSO_4_ and 0.1 mM CaCl_2_. As carbon and energy source, either 27.8 mM [0.5% w/v] glucose, 6.47 mM [0.2% w/v] 5-N-acetyl neuraminic acid (Neu5Ac), or 6.15 mM [0.1% w/v] N-glycolylneuraminic acid (Neu5Gc) (all purchased from Sigma-Aldrich, Taufkirchen, Germany) was added. If appropriate, the following antibiotics were used: ampicillin sodium salt (150 μg/ml) or kanamycin (50 μg/ml). For solid media, 1.5% agar (w/v) was added. For all growth experiments, bacterial strains were grown in LB medium overnight at 37°C, washed twice in PBS and then adjusted to an optical density at 600 nm (OD_600_) of 0.005 in the desired liquid growth medium, or streaked on agar plates. Growth curves were obtained from bacterial cultures incubated at 37°C with gentle agitation in 96-well microtitre plates containing 200 μl medium. The OD_600_ was measured by an automatic reader (Epoch2T; BioTek, Bad Friedrichshall, Germany) at appropriate time intervals as indicated.

## Results

### Data collection

After WGS quality control, 14 isolates were excluded because of poor quality sequences. One additional isolate was excluded since this isolate was identified as *Escherichia marmotae* (formerly cryptic clade V)(43,44), a species commonly mistaken for *E. coli.* Our final collection comprised 1198 *E. coli* whole-genome sequences with metadata (Table S1), which also contained 8 cryptic clade I isolates, which were included as *E. coli* based on the recommended species cut-off of 95-96% average nucleotide identity(43). Our collection consisted of 22.1% (n=265) cattle, 28.1% (n=337) chicken, 27.3% (n=327) human, 20.3% (n=240) pigs and 2.4% (n=29) wild boar isolates (Fig. S1A). Fifty-one percent (n=612), 19.4% (n=233), 14.5% (n=174) and 14.9% (n=179) of these isolates were from Germany, Spain, the UK, and Vietnam, respectively (Fig. S1A). Chicken isolates were from all four countries, human isolates from Germany, the UK and Vietnam, pig isolates from Germany, Spain and Vietnam, cattle isolates from Germany and Spain and only Spain provided wild boar isolates. In total, 35.5% (n=426) of the isolates were from hosts with reported disease, whereas 62.0% (n=743) were from hosts without reported disease, while host health status was unknown for the wild boar isolates (2.4%, n=29). Of the 1198 isolates analyzed, 1140 were grouped into 358 different STs, and 58 could not be assigned to any known ST. The population structure of the collection closely resembles that of the ECOR collection(45), indicating that it represents most of the known diversity of *E. coli sensu stricto* (Fig. S2).

### Pan-genome analysis

The *pan*-genome of the 1198 *E. coli* isolates consisted of 77130 genes, of which 1956 genes belonged to the core genome (i.e., present in at least 99% of the isolates). The population structure of the collection based on core genome single-nucleotide polymorphisms (SNPs) was defined using Bayesian analysis of population structure (BAPS), which assigns isolates to discrete clusters. Most of the isolates were assigned to phylogroups B1 (n=366, 30.55%), A (n=313, 26.12%) and B2 (n=213, 17.77%). The remaining isolates were distributed among phylogroups D (n=97, 8.09%), E (n=55, 4.59%), G (n=49, 4.09%), F (n=35, 2.92%), C (n=60, 5.0%), and clade I (n=8, 0.6%). A comparison of phylogenetic clusters, phylogroups, country, host, and a maximum likelihood (ML) tree based on 110920 core-genome SNPs is shown in Fig 1. The χ^2^-tests for independence revealed a positive correlation between host status and phylogenetic clusters (at *p* < 2.26e^-16^, df=52) and between phylogroups and hosts (*p*<2.2e^-16^, df=32). This indicates that specific phylogenetic clusters (Fig. S1 B&C) and phylogroups, such as B1 (cattle), A (pig), B2 (human and chicken), and G (chicken) were enriched within different hosts in our collection (Fig. S1D).

**Fig 1:**
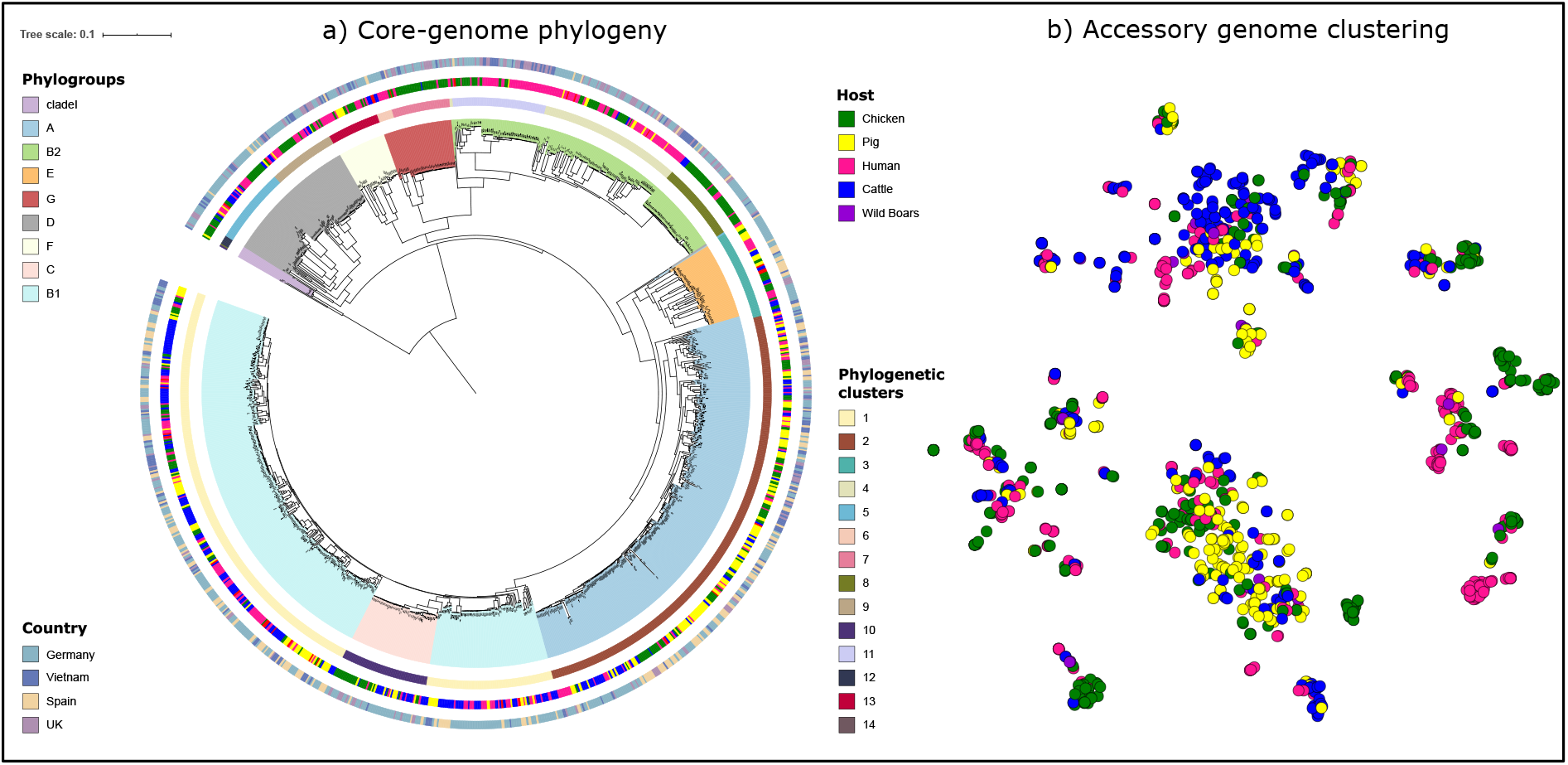
Distribution of 1198 isolates with host species by a) core-genome phylogeny and b) clustering based on accessory gene content (right). Clades on the phylogeny represent phylogroups, inner-ring represents phylogenetic clusters, middle-ring represents host-species, and outer ring indicates the geographical region.

Clustering of isolates based both on core gene alignment and on accessory gene profile appeared to be correlated with phylogroups. The interactive visualization of data is also available on Microreact (https://microreact.org/project/ouDOdcFxc). A minimum spanning tree was built on the allelic profiles of 358 (n=1,140 isolates) known STs and 58 isolates belonging to unknown STs using GrapeTree(46) along with the host distribution (Fig. S3). Several sequence types, of which at least ten isolates were available, appeared to be linked with certain host species. ST33 (n= 10/10, 10 human isolates out of all 10 isolates), ST73 (n=11/17), ST131 (n=37/42) and ST1193 (n=12/12) were associated with a human host. ST131 was also found in chickens (n=4/42) and pigs (n=1/42) in this collection. ST23 (n=18/22), ST95 (n=25/31), ST115 (n=11/11), ST117 (n=30/33), ST140 (n=19/20) and ST752 (n=29/30) were associated with the chicken host.

### GWAS

The genome-wide association analysis was performed on 1169 *E. coli* isolates from cattle, chickens, humans, and pigs. The 29 wild boar isolates were excluded because of their small group size. Genome-wide association analysis revealed the positive association (β>0) of 27,854, 16,164, and 69,307 *k-mers* with *E. coli* isolates from humans, cattle, and chickens at a likelihood ratio test*p-value* less than 1.87×10^-9^, 2.16×10^-9^, and 1.9×10^-9^ respectively (reported as “lrt-pvalue”). There were no *k-mers* significantly associated with the pig host. The significant *k-mers* accounted for 426, 179, and 915 bacterial genes associated with isolation from human, cattle, and chicken hosts, respectively (Fig 2 and Table S3). An arbitrary cut-off of at least 10 *k-mers* mapped per gene was chosen to select genes for *in silico* functional characterization as well as COG assignment using Blast2GO(38) (Table S4) and CD-search(39,40) (Fig. S4).

**Fig 2:**
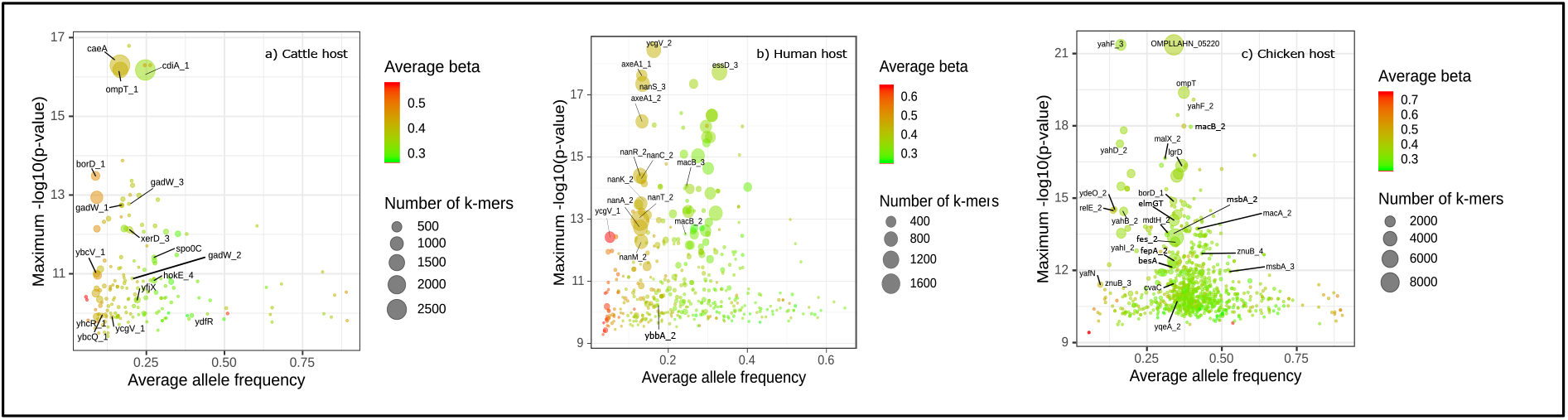
Plots representing the *E. coli* genes or gene variants associated with the a) Cattle host, b) Human host, and c) Chicken host. The bubble size represents the number of k-mers mapped to a specific gene, and the color gradient represents the effect size (β).

### Association of novel nan genes with human host

GWAS revealed a strong association of nine contiguous genes, assigned to the group of *nan* genes with the human host (Fig 2b). Seven of these genes were annotated *in silico* as *nan* genes (Fig 3a) and the remaining two genes were annotated as being similar to *axeA1* of *Prevotella ruminicola* ATCC 19189 (Uniprot accession D5EV35). However, the amino acid sequences of the products of these *axeA1-like* genes only shared 19-20% similarity with AxeA1. Further investigation with EggNOG and CD search revealed an acetylesterase/lipase-encoding region (COG0657) in both genes and confirmed *nan* gene annotations. Previous evidence and the genomic location (i.e., between the *nan* genes; Fig 3a) suggest that these genes encode potential acetylesterases and may be analogous to sialyl esterases (NanS)(47). Hence, these nine novel *nan* genes are collectively termed “human-associated *nan* gene cluster (*nan*-9)” (Fig 3a).

**Fig 3:**
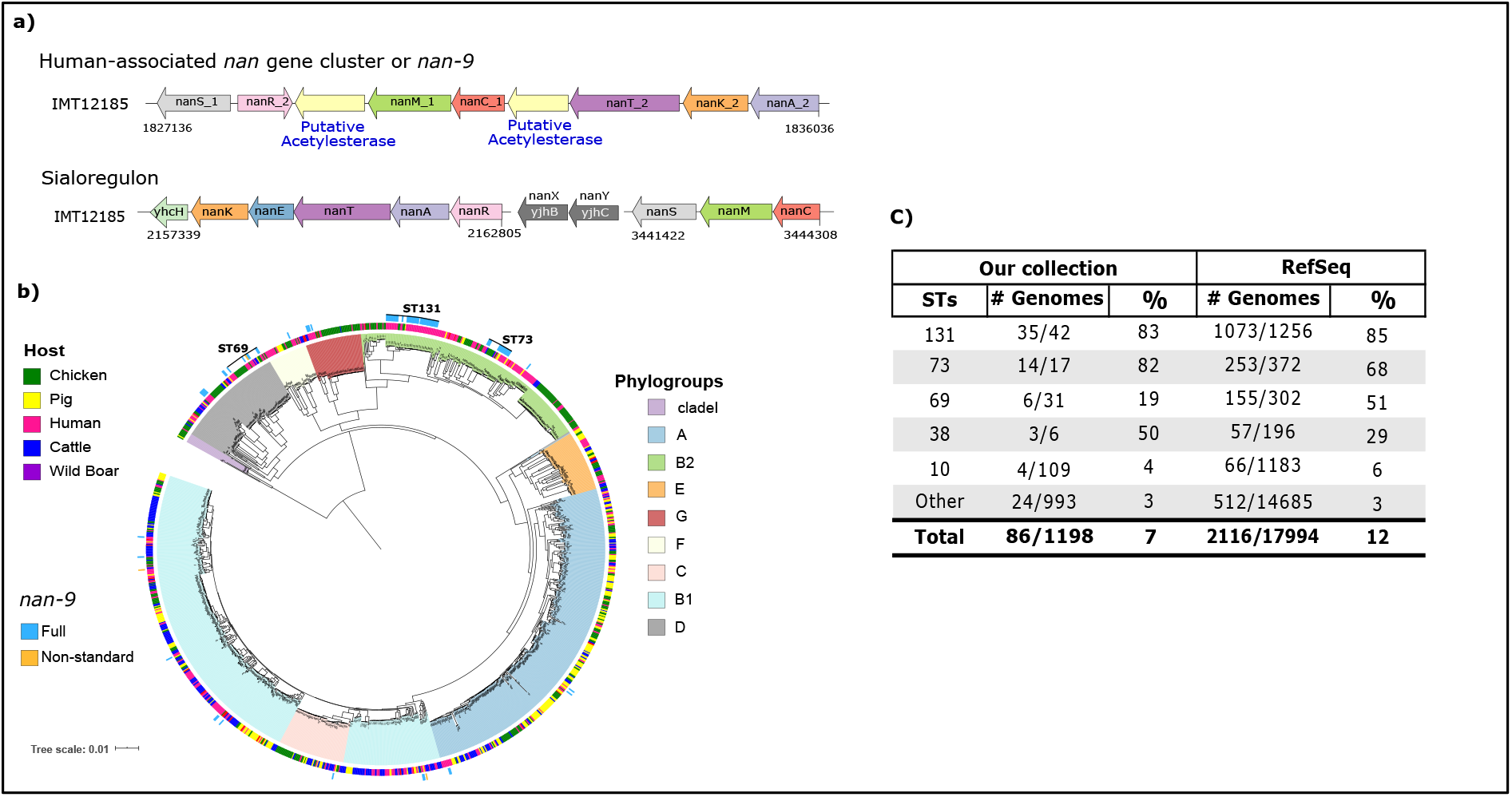
a) Genetic architecture of the human-associated *nan* gene cluster *(nan-9)* and the sialoregulon on the complete genome of the strain IMT12185. The strain lacks the *nanXY* genes of the sialoregulon. b) Distribution of the *nan-9* cluster on core-genome phylogeny marked with STs with higher prevalence. c) The table indicates the prevalence of the *nan-9* gene cluster in different STs in our collection and in the RefSeq *E. coli* genomes.

Distinct *nan* genes are present in *E. coli* and are also known as the sialoregulon (*nanRATEK-yhcH, nanXY[yjhBC],* and *nanCMS;* Fig 3a)(48). The sialoregulon is known to be involved in metabolism of sialic acids(49–51), a diverse group of nine-carbon sugars, abundant in the glycocalyx of many animal tissues(52,53). Sialic acids present on mucin proteins in the human gut are an essential energy source for many intestinal bacteria(54). The proteins encoded by the seven genes of *nan-9* (i.e. *nanAKTCMRS)* share 45-64% similarity with the corresponding *nan* genes of the sialoregulon in *E. coli* or the recently described phage-encoded *nanS-p* genes of enterohemorrhagic *E. coli(55).* Both the human-associated *nan* gene cluster and the sialoregulon are located on the bacterial chromosome. The human-associated *nan* gene cluster was found in 7% of our isolate collection, whereas the genes comprising the sialoregulon were more common. In our collection, *nanXY* was identified in ~15% of isolates, *nanCMS* in ~93% of isolates, whilst *nanRATEKyhcH* was found in almost all (>99%) isolates.

The *nan-9* cluster was detected in 86 isolates, mainly from phylogroups B2 and D (Fig 3b) and predominantly in isolates belonging to ST131, ST73, and ST69, both in our collection as well as across 17,994 RefSeq *E. coli* genomes (Fig 3c). The order and orientation of genes in the human-associated *nan* gene cluster were found to be identical in 82 out of 86 isolates (Fig. S5). In 63 isolates, insertion sequence (IS) 682 was found upstream, and in 23 isolates, IS2 was found downstream of this novel gene cluster (Fig. S5).

To further explore the function of the human-associated *nan-9* gene cluster, the entire cluster was knocked-out from strain IMT12185 (ST131), yielding strain IMT12185Δ*nan-9*. For comparison, an additional mutant, which lacked the *nanRATEK* locus from the sialoregulon (ΓMT12185Δ*nanRATEK*) was constructed from wild-type IMT12185. Correct gene deletion in both mutants was confirmed through WGS. No significant differences in carbon utilization and chemical sensitivity were observed between wild-type strain IMT12185 and its mutant IMT12185Δ*nan-9* in Biolog phenotyping array experiments (PM1 and Gen III MicroPlates).

Deletion mutant IMT12185Δ*nan-9* was grown in MM with 0.2% 5-N-acetylneuraminic acid (Neu5Ac) or with 0.1% N-glycolylneuraminic acid (Neu5Gc) as sole carbon and energy source. Neu5Ac is the most common sialic acid of the glycocalyx of both humans and other mammals, whereas Neu5Gc is absent in humans. In the presence of Neu5Ac, mutant IMT12185Δ*nan-9* grew to a maximal OD_600_ of 1.34 comparable to that of parental strain IMT12185 (OD_600_ = 1.37). However, the mutant exhibited a delayed growth start of approximately three hours (Fig 4a). When Neu5Gc was offered as substrate, the mutant not only showed a similar growth start retardation, but also a slower growth rate and a lower maximal OD_600_ (1.31) in comparison with strain IMT12185 (OD_600_ = 1.43) (Fig 4b). Both Neu5Ac and Neu5Gc are degraded by the enzymatic activities of the enzymes NanRATEK, of which four, namely NanRATK, are encoded by redundant genes located on the determinants *nanRATEK* and *nan-9.* Deletion mutant IMT12185*ΔnanRATEK* was unable to grow with Neu5Ac (Fig 4c), demonstrating that *nan-9* alone is not sufficient for sialic acid degradation, probably due to a lack of *nanE* in the *nan-9* gene cluster. To exclude a pleiotropic effect of the *nan-9* deletion, parental strain IMT12185 and its mutant IMT12185Δ*nan-9* were grown in LB medium. No significant difference was observed between the two growth curves (Fig 4d). These data demonstrate that the *nan-9* determinant of strain IMT12185 is biologically functional and contributes to the degradation of the sialic acids Neu5Ac and Neu5Gc.

**Fig 4:**
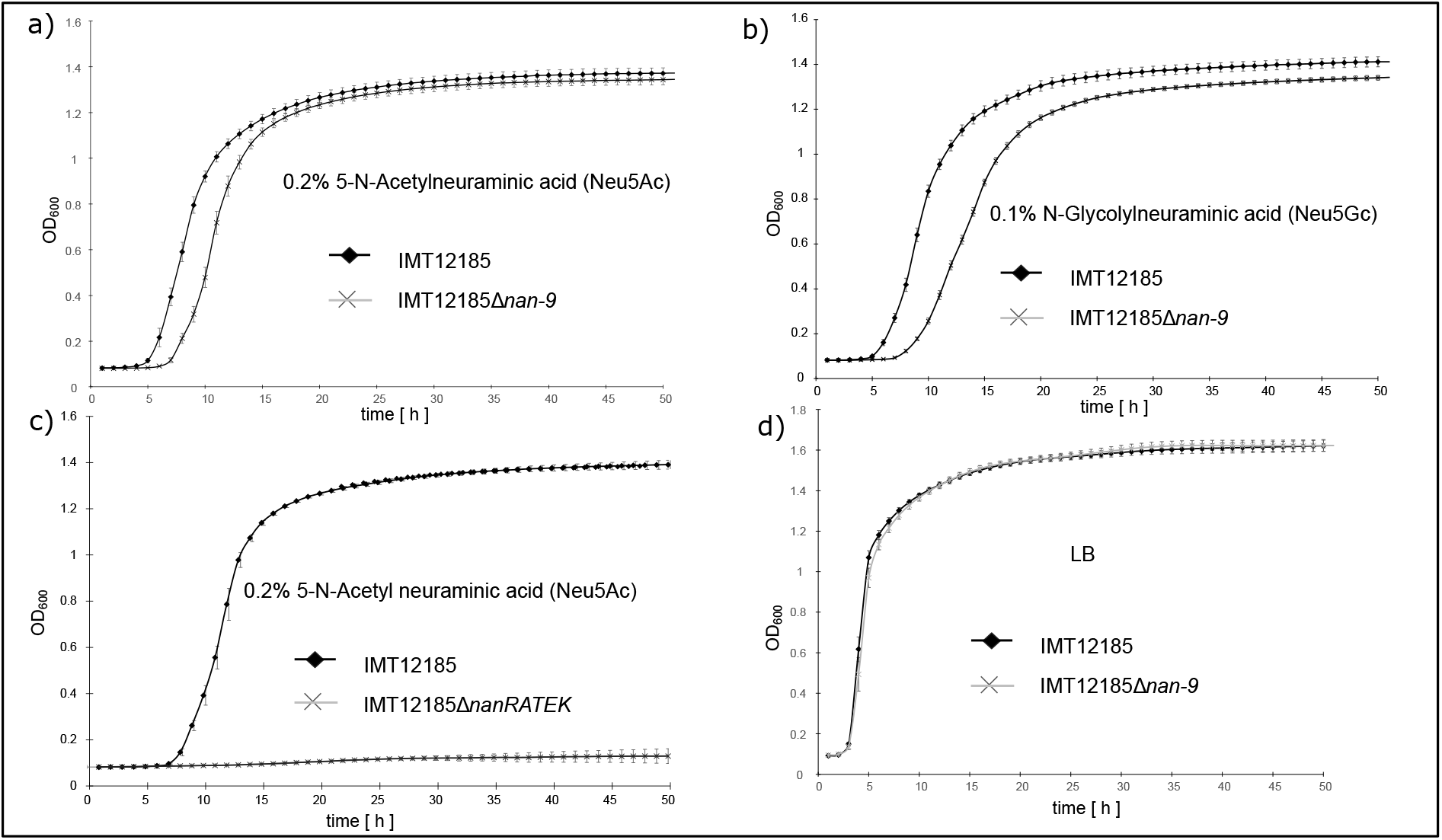
Growth curves of *E. coli* IMT12185 and its mutant derivatives in various media. a) Growth of IMT12185 and IMT12185△nan-9 in M9 minimal medium with 0.2% 5-N-Acetylneuraminic acid (Neu5Ac) b) Growth of IM12185 and IMT12185△nan-9 in M9 minimal medium with 0.1 5-N-Glycolylneuraminic acid (Neu5Gc) c) Growth of IMT12185 and IMT12185△nanRATEK in M9 minimal medium with 0.2% 5-N-Acetylneuraminic acid (Neu5Ac) d) Growth of IMT12185 and IMT12185△nan-9 in lysogeny broth (LB).

### Other genes associated with the human host

Several other genes associated with the human host were identified in the GWAS analysis, such as the *sat* gene encoding a serine protease autotransporter vacuolating toxin (Fig 2b)(56). This gene was detected in 22.9% (n=75/327) of the human isolates in our collection and in only 0.59% (n=5/891) of the strains isolated from other hosts (Table S5). This gene was mainly detected in isolates belonging to specific lineages such as ST131, ST1193, and ST73 (Table S5). In addition, we found an association with two distinct homologs of the *macB* gene that encodes an ABC transporter(57) and is involved in many diverse processes, such as resistance to macrolides(58), lipoprotein trafficking(59), and cell division(60).

### Association of distinct Omptins with the cattle and chicken hosts

We detected homologs of the *ompT* (encoding outer-membrane protease VII) gene, a member of the omptin family of proteases, in our dataset (Fig 2a & Fig 2c). Two homologs, *ompP* (UniProt accession P34210, sharing 70% amino acid identity with OmpT) and *arlC* (also referred to as *ompTp,* UniProt accession Q3L7I1, sharing 74% amino acid identity with OmpT), were found to be associated with the cattle and chicken hosts, respectively (Fig 5). In our collection, *ompP* was predominant in phylogroup B1 (n=68), whereas *arlC* was found in distinct phylogroups (such as B2, B1 and G) (Fig 5) and in isolates belonging to ST95 and ST117 (Table S6). A similar association was observed in 17,994 public *E. coli* genomes from RefSeq (Table S6). Previous studies have reported an increased prevalence of *arlC* (erroneously reported there as *ompT*) in a cluster of uropathogenic *E. coli* (UPEC) and avian pathogenic *E. coli* (APEC) classified as ST95(61). Notably, *arlC* is associated with increased degradation of antimicrobial peptides (AMPs) in UPEC isolates(62). OmpP is also able to degrade AMPs and displays a AMP cleavage specificity different from that of OmpT(63).

**Fig 5:**
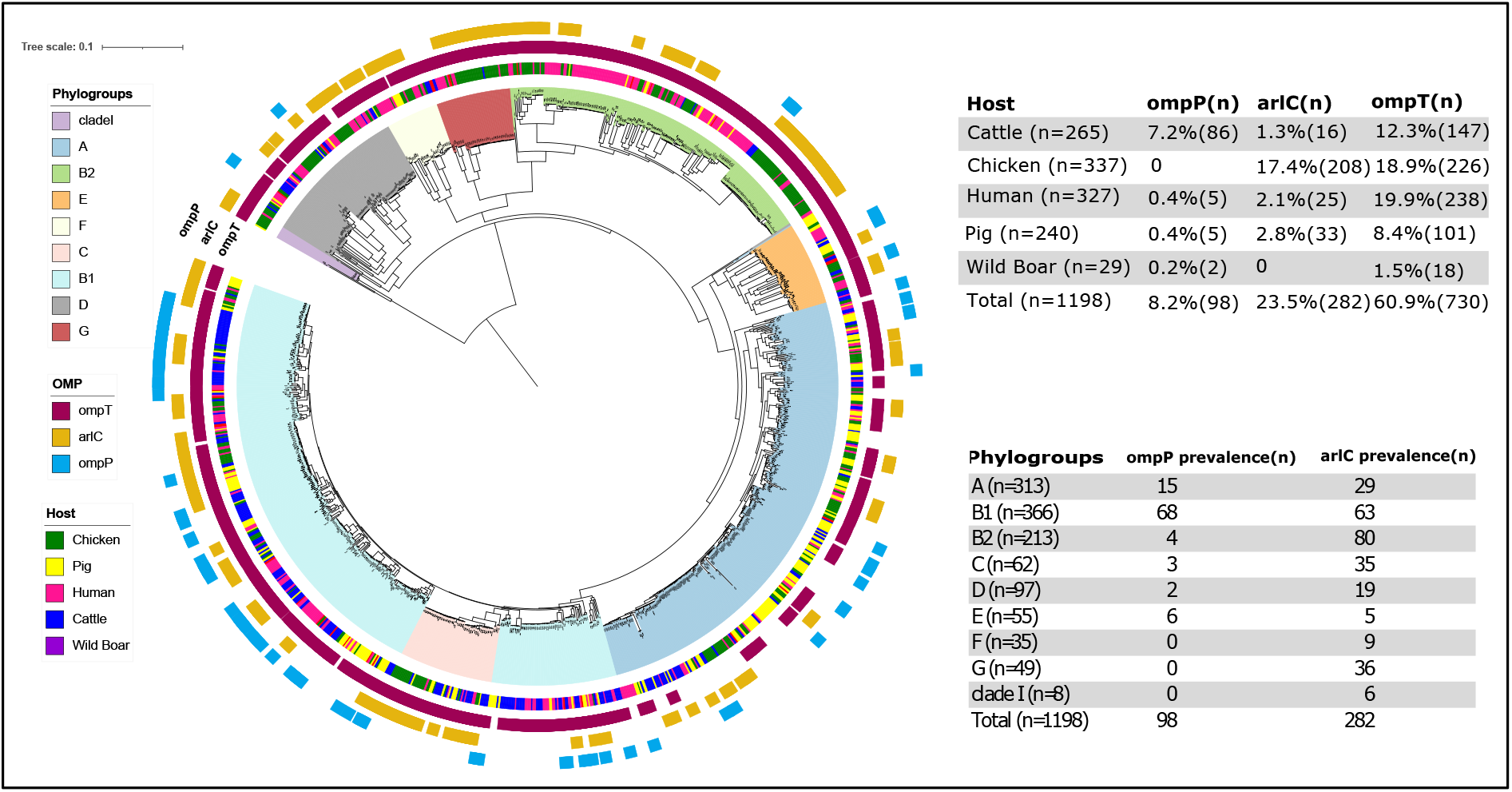
Distribution of *ompP, arlC,* and *ompT* genes in phylogroups and host across the phylogeny and their estimated prevalence.

### Association of genes involved in metal acquisition with the chicken host

GWAS analysis revealed an association of the *iroBCDEN* gene cluster (C) with the chicken host, but not with other host species included in this study. The prevalence of the *iro* gene cluster was 24.3% (n=291/1198) in our collection, of which 61.5% (n=179/291) were from the chicken host. The gene cluster was found in different STs and with higher prevalence in STs such as ST117, ST95, ST23, and ST140 (Table S7). The chromosomal *iroBCDEN* gene cluster was first described in *Salmonella enterica* and is involved in uptake of catecholate-type siderophores, high-affinity iron-chelating molecules contributing to bacterial survival during infection by sequestering iron(64). In *E. coli,* this gene cluster has mainly been described in uropathogenic (UPEC) and avian pathogenic *E. coli* (APEC) and is regarded as a virulence factor(65). The cluster has been reported on a chromosomal pathogenicity island, although in ExPEC, the cluster can also be located on ColV or ColBM virulence plasmids(66,67). In addition, homologs of genes involved in zinc catabolism *(znuB)* and iron metabolism *(fes)* were found to be associated with the chicken host (Fig 2c).

## Discussion

*Escherichia coli* can colonize many different ecological niches in a diverse range of host species, ranging from a commensal lifestyle to intra-or extra-intestinal infections. Presence of certain adhesin and other virulence-associated genes is well known to correlate with the relative ability of *E. coli* strains to colonize the intestinal tract of certain hosts (e.g., *ecp* for humans(68), F9 fimbriae and H7 flagellae for cattle(69,70) or Stg fimbriae for chickens(71)). Variations in host adaptation levels and their molecular basis in *E. coli* strains presumptively realizing a commensal-like lifestyle in the reservoir host are rarely described and poorly understood as of yet(72). Commensal *E. coli* strains may be carriers of AMR and a source of mobile genetic elements conferring AMR to other bacteria including pathogenic strains in a shared microbiome, e.g. in the intestinal tract of animals including humans. We therefore collated an extensive and diverse dataset to identify genetic determinants of *E. coli* host adaptation. We observed significant enrichment of specific hosts within some phylogroups and STs in our collection. Furthermore, we unveiled correlations between the likelihood of genetically related isolates having been isolated from a certain host with the possession of distinctive genetic traits. Some of these traits, e.g. the *iroBCDEN* gene cluster, have been linked to *E. coli* and *Salmonella* virulence before, while others, in particular the human-associated *nan* gene cluster, are novel traits and have not been implicated in the infection and colonization process of *E. coli.* Of note, the latter gene cluster encodes for metabolic properties which have received little attention in bacterial infectious disease research. Specific metabolic properties have been linked to the relative ability of Shiga toxin-encoding *E. coli* (STEC) to asymptomatically colonize cattle, their reservoir host(73). Unraveling the nutrient and energy flows in the complex interplay of intestinal bacteria, the surrounding microbiome and the host may open novel avenues to control the persistence and transmission of pathogenic and/or antimicrobial resistant bacteria(74).

We employed a *k-mer* based bacterial GWAS, applied in previous studies to associate multiple types of genetic variation with phenotypes(75,76). In our study, we were able to associate a phenotype (i.e., isolates obtained from a certain host species) with the presence of specific genes, but not with sequence variation at the level of single nucleotide polymorphisms between genes. This lack of associations found at the SNP level could possibly be explained by the fact that through our filtering approach to prevent false positive hits, we might have excluded *k-mers* that captured host-associated SNP variation. Secondly, it might be possible that since *E. coli* is genetically diverse, host-associated SNP variation is challenging to capture between unrelated strains. Finally, the absence of host-associated SNPs might be a biological observation, indicating that colonization of particular hosts is determined by gene presence or absence rather than minimal genetic variation within genetic elements. However, we were able to confirm previously published host associations, indicating the validity of our approach. For example, carriage of the salmochelin operon encoded by *iroBCDEN* and involved in iron metabolism was previously identified as associated with increased ability of *E. coli* strains to colonize chickens(65,77).

In addition to *iroBCDEN,* we found an association of omptin proteins (OmpP and ArlC) with chickens and cattle as hosts, respectively. Earlier studies using UPEC strains had demonstrated that these proteins are associated with cleavage and inactivation of cationic antimicrobial peptides (AMPs)(62). Because AMPs are secreted as part of the host’s innate immune response(78–80), these proteins may play a vital role in colonization. AMPs are also increasingly used as alternatives to antimicrobial agents in animal farming(81–83), further investigation into the contribution of these Omp variants to host colonization as well as to resistance to exogenous AMPs is warranted.

We did not identify any significant associations of *k-mers* with the pig host. Bacterial colonization of the porcine intestine by edema-disease *E. coli* (EDEC) is mediated by the ability of these bacteria to adhere to villous epithelial cells via their cytoadhesive F18 fimbriae(84). The expression of receptors for these fimbriae on the apical enterocyte surface is inherited as a dominant trait among pigs and determines susceptibility to diseases caused by F18-fimbriated pathogenic *E. coli*(85). Enterotoxigenic *E. coli* (ETEC) express F4 or F5 fimbriae with similar consequences(86). However, we found only three, four and six isolates harbouring genes for F4, F5 and F18 fimbriae, respectively. Thus, we might not have had all *E. coli* pathovars associated with pig host sufficiently present in our collection, although we did observe an association between phylogroup A and pig colonization. An alternative reason might be that the association between phylogroup A and pig colonization complicated the identification of statistically significant *k-mers*. GWAS corrects for population structure, which means that if there is a strong association between lineage and phenotype, the genes harbored by that lineage will not be reported as having a strong association with the phenotype under study(87).

We identified a novel human host-associated *nan* gene cluster, distinct from the previously reported sialic acid (Sia) metabolic operon *(nanRATEK-yhcH, nanXY,* and *nanCMS)(48).* This novel cluster is conserved and abundant in ExPEC lineages, such as ST131, ST73, and ST69. The gene cluster is flanked by insertion sequences which might play a role in the horizontal exchange between different *E. coli* lineages. Knock-out *in vitro* studies indicated that this novel *nan-9* gene cluster contributes to catabolism of the sialic acids Neu5Ac and Neu5Gc, although it cannot replace the function of the *nanRATEK*locus which is abundant in *E. coli.* Hence, we hypothesize that *E. coli* harboring the *nan-9* gene cluster have an evolutionary advantage through either more efficient access to sialic acids or through access to more diverse sialic acids. The genes annotated as acetylxylan esterases are expected to represent novel sialyl esterases, as known sialyl esterases (*nanS* variants) have previously been mistaken for acetylxylan esterases(47). Additional sialyl esterases – possibly with alternative deacetylation specificity – might provide a more efficient catabolism of acetylated sialic acids. Future studies should investigate the role of the human-associated *nan-9* gene cluster in the catabolism of differentially acetylated sialic acids and their relevance for the human host.

Approximately one-third of the isolates in our dataset were obtained from diseased hosts, while the remaining isolates were from healthy hosts. Many of the isolates in our dataset that originate from healthy hosts belong to ExPEC lineages which are typically considered to be pathogenic. In fact, the locus most strongly associated with the human host, the *nan-9* gene cluster, is abundant in ExPEC lineages. This does not necessarily mean that the *nan-9* gene cluster is associated with pathogenicity. In fact, this observation primarily supports the notion that these pathogenic *E. coli* are highly efficient colonizers of the human intestine(72). Based on our results, we hypothesize that the human-associated *nan-9* gene cluster is one of the factors driving the adaptation of ExPEC to the human intestine.

Finally, we observed an association between the *sat* gene and human host colonization. Sat contributes to the pathogenicity of *E. coli* in the urinary tract(56). The high prevalence of *sat* in previously studied *E. coli* isolates from the feces of healthy individuals suggests it may not act as a virulence factor in the human gut(88). However, in our isolate collection, the *sat* gene was found in *E. coli* strains belonging to phylogroups A, B2, D, and F, which had been isolated from both healthy and diseased hosts (Table S5). Understanding the role of Sat in the colonization and adaptation of *E. coli* in healthy humans warrants further investigation.

## Conclusion

Our study identified several distinct genetic determinants that may influence *E. coli* adaptation to different host species and provide an adaptive advantage. These findings are important as they aid the better understanding of the potential outcome of transmission events of *E. coli* between host species. This is particularly relevant for the control of the spread of antimicrobial resistant commensal and zoonotic *E. coli* strains within and across human and animal populations. The data generated here can also be used in risk analysis and for diagnostic and monitoring purposes. More importantly, our study identified biological processes, including sialic acid catabolism, that should be investigated in more detail to better understand *E. coli* host adaptation.

## Data availability

The raw-reads of the 1090 *E. coli* isolates sequenced in this study were submitted to NCBI SRA with the Bioproject accession number PRJNA739205 and the SRA accession of 108 isolates, that were taken from other studies, were provided in supplement table S1.

## Acknowledgements

The HECTOR research project was supported under the framework of the JPIAMR – Joint Programming Initiative on Antimicrobial Resistance – through the 3^rd^ joint call, thanks to the generous funding by the Netherlands Organisation for Health Research and Development (ZonMw, grant number 547001012), the Federal Ministry of Education and Research (BMBF/DLR grant numbers 01KI1703A, 01KI1703C and 01KI1703B), the State Research Agency (AEI) of the Ministry of Science, Innovation and Universities (MINECO, grant number PCIN-2016-096), and the Medical Research Council (MRC, grant number MR/R002762/1).

