## Supplementary figures for "Genome-wide association reveals host-specific genomic traits in *Escherichia coli*"

**Fig. S1:** Distribution of 1,198 isolates and enrichment analysis: **A)** The plot represents the proportion of *E. coli* strains isolated from hosts in four countries. The number above each plot indicates the total number of isolates per host. **B)** Proportion of *E. coli* strains obtained from the five host species contributing to 14 phylogenetic clusters. The number on top of each bar represents the total number of isolates per cluster. **C)** Phylogenetic clusters enriched with a different host (Pearson residual > 0 represents positive correlation indicating the enrichment of certain host-species in distinct clusters at *p-value* <2.22e$^{-16}$). **D)** The proportion of *E. coli* isolates from different host species contributing to different phylogroups.

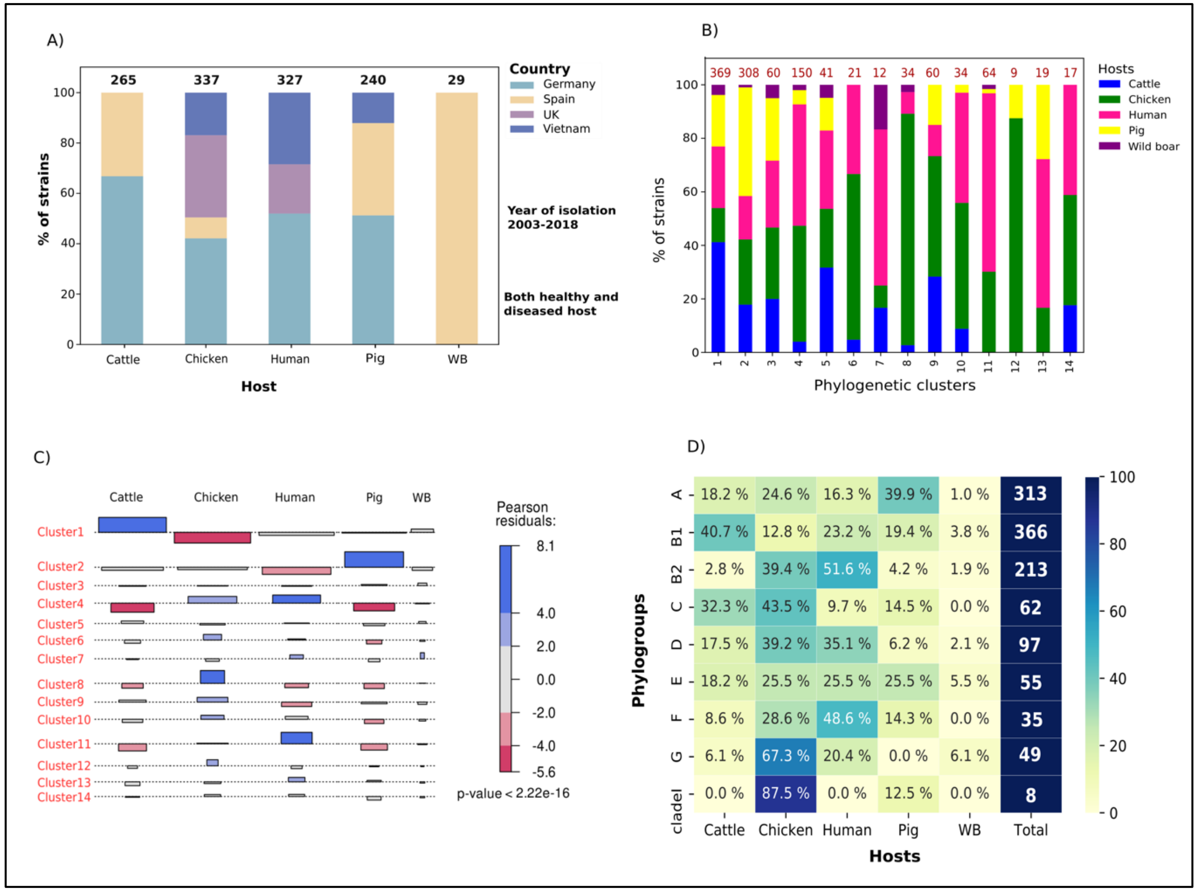

**Fig. S2:** Core genome phylogeny of *E. coli* isolates of our collection (n=1,198) and Reference strains (n=146) from the ECOR collection, RefSeq and cryptic clades annotated with their phylogroups (phylogroups were determined by ClermonTyper v. 1.3).

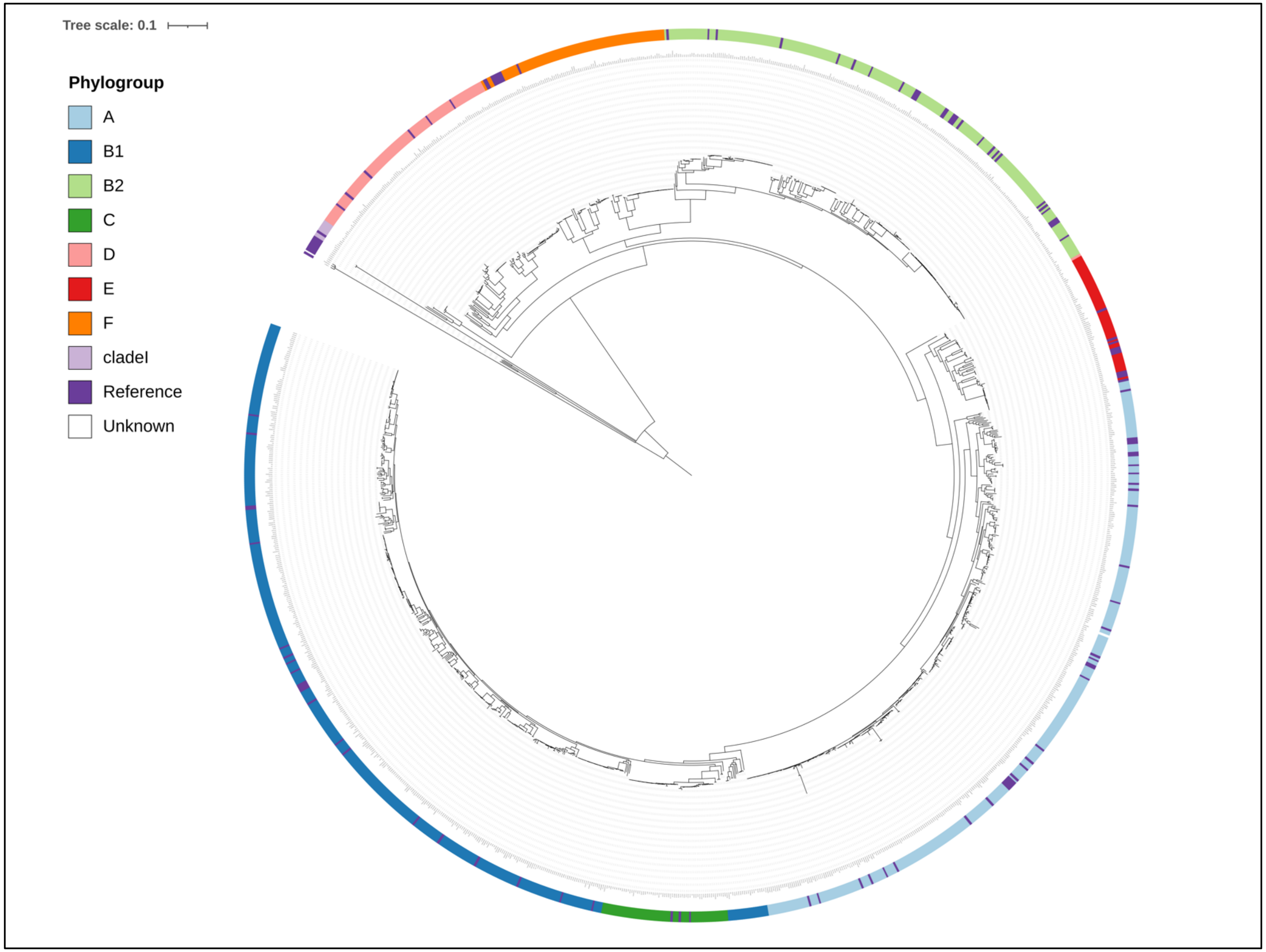

**Fig. S3:** Minimum-spanning tree of MLST profiles of 1,198 *E. coli* isolates. Left: the number of isolates constituting an ST; Right: the proportion of hosts in each ST.

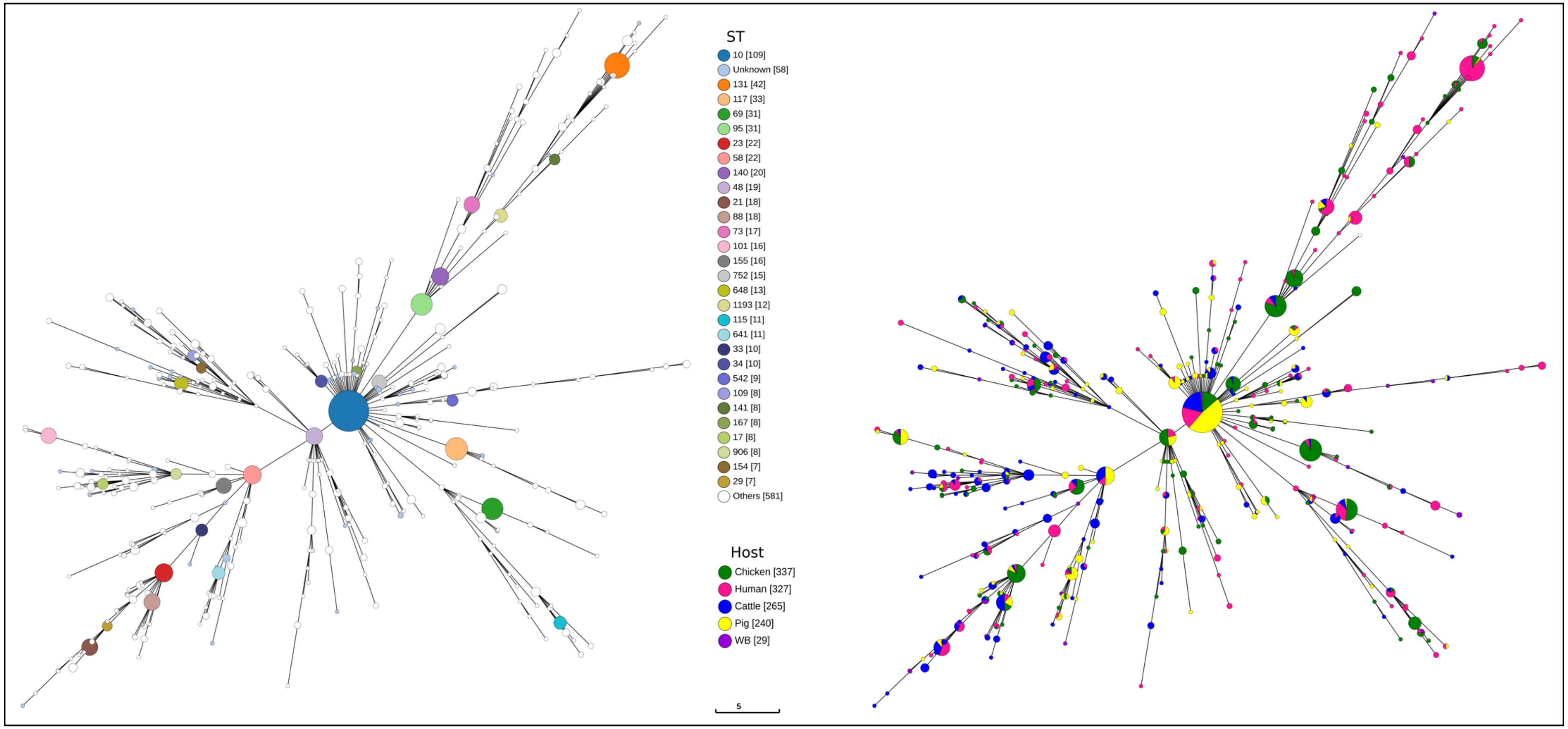

**Fig. S4:** COG classification of genes or gene variants associated with cattle, chickens, and humans. Y-axis: Indicates the molecular function, and the x-axis indicates the number of genes in each functional class from each host.

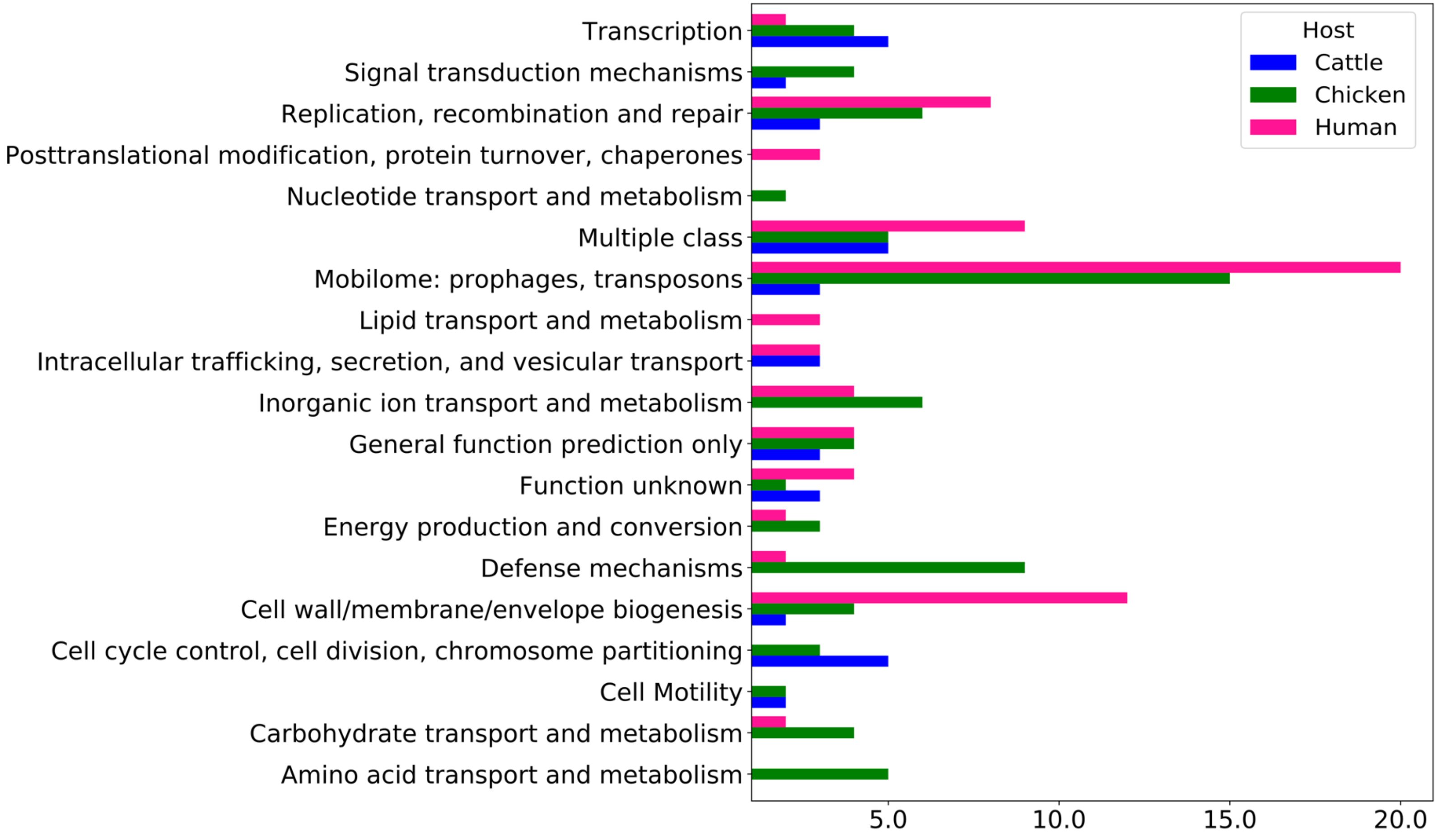

**Fig. S5:** Genetic surroundings of the human-associated *nan* gene cluster in the genomes of all isolates in which it was identified in.

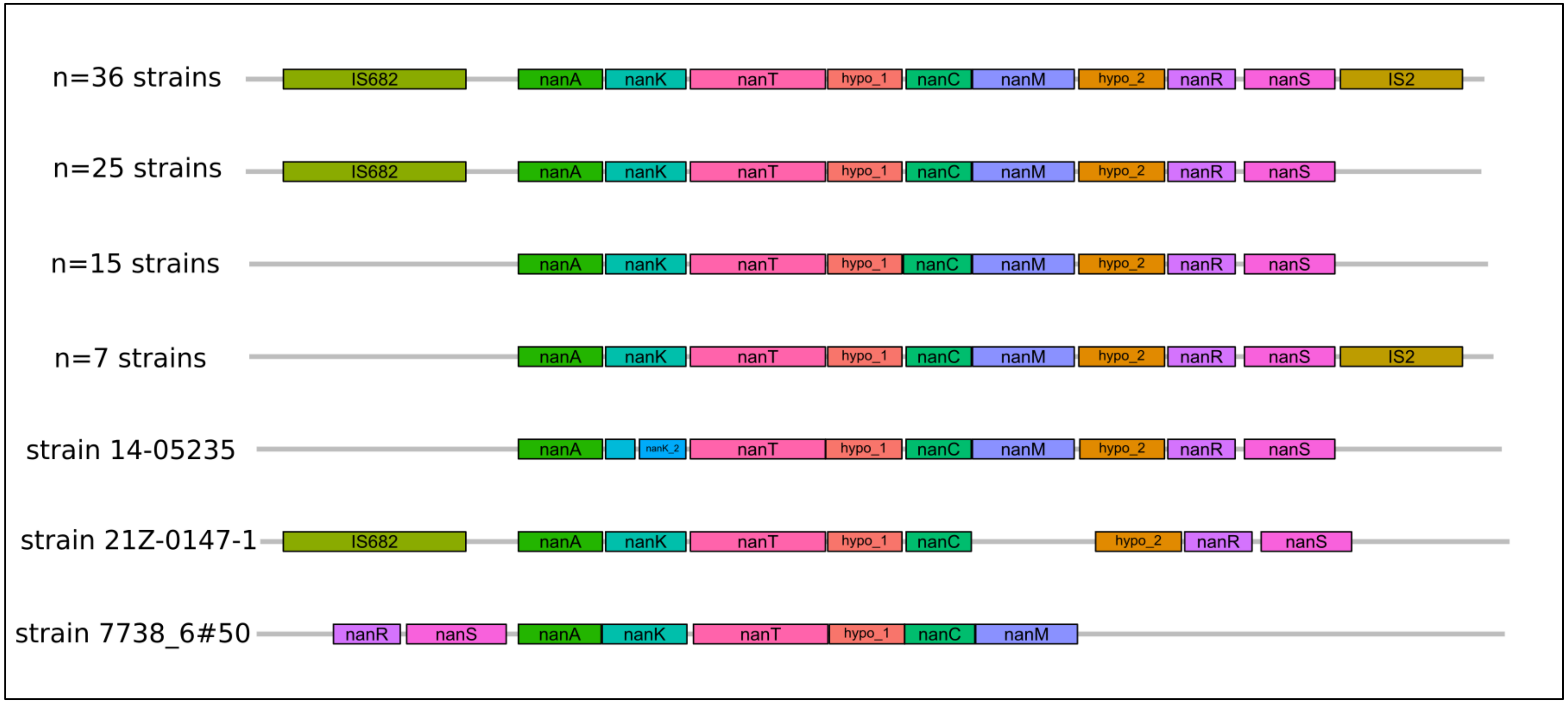
